# A Rare Deletion in SARS-CoV-2 ORF6 Dramatically Alters the Predicted Three-Dimensional Structure of the Resultant Protein

**DOI:** 10.1101/2020.06.09.134460

**Authors:** Marco A. Riojas, Andrew M. Frank, Nikhita P. Puthuveetil, Beth Flores, Michael Parker, Stephen P. King, Malik Peiris, Daniel K. W. Chu, Briana Benton, Rebecca Bradford, Manzour Hernando Hazbón, Sujatha Rashid

## Abstract

The function of the SARS-CoV-2 accessory protein p6, encoded by ORF6, is not fully known. Based upon its similarity to p6 from SARS-CoV, it may play a similar role, namely as an antagonist of type I interferon (IFN) signaling. Here we report the sequencing of a SARS-CoV-2 strain passaged six times after original isolation from a clinical patient in Hong Kong. The genome sequence shows a 27 nt in-frame deletion (Δ27,264-27,290) within ORF6, predicted to result in a 9 aa deletion (`ΔFKVSIWNLD`) from the central portion of p6. This deletion is predicted to result in a dramatic alteration in the three-dimensional structure of the resultant protein (p6^Δ22-30^), possibly with significant functional implications. Analysis of the original clinical sample indicates that the deletion was not present, while sequencing of subsequent passages of the strain identifies the deletion as a majority variant. This suggests that the deletion originated *ab initio* during passaging and subsequently propagated into the majority, possibly due to the removal of selective pressure through the IFN-deficient Vero E6 cell line. The specific function of the SARS-CoV-2 p6 N-terminus, if any, has not yet been determined. However, this deletion is predicted to cause a shift from N-endo to N-ecto in the transmembrane localization of the SARS-CoV-2 p6^Δ22-30^ N-terminus, possibly leading to the ablation of its native function.

## Introduction

In late 2019, a previously unknown coronavirus emerged and began causing severe human respiratory infections centered primarily in Wuhan, Hubei Province, China. The initial infections reported included atypical pneumonia, fever, dyspnea, and dry cough as symptoms.^1^ The virus was initially referred to by the provisional name 2019 novel coronavirus (2019-nCoV) but was later officially designated “severe acute respiratory syndrome-related coronavirus 2” (SARS-CoV-2) by the International Committee on Taxonomy of Viruses (ICTV)^2^, and the disease it causes was named COVID-19 by the World Health Organization (WHO)^3^. At the time of writing, approximately six months after it was first identified, the outbreak of COVID-19 has spread to virtually every country in the world, infecting over 7.2 million people and killing over 410,000 people thus far.^4,5^ In response to this pandemic, the global scientific community has been intensely researching the virus in an effort to develop diagnostics and countermeasures.

Since January 2020, the Biodefense and Emerging Infections Resources (BEI Resources) has been building a comprehensive catalog of strains and related reagents to support organizations performing COVID-19 research and development. BEI Resources was established by the National Institute of Allergy and Infectious Diseases (NIAID) to provide reagents, tools and information for studying Category A, B, and C priority pathogens, emerging infectious disease agents, non-pathogenic microbes and other microbiological materials of relevance to the research community. On February 4th, 2020, BEI Resources acquired its first SARS-CoV-2 strain, which was isolated from the first reported US patient, from the Centers for Disease Control (CDC) and immediately produced, characterized, and began distributing this isolate on February 6th, 2020. To date, BEI Resources has provided more than 7,700 vials of SARS-CoV-2 and related coronavirus strains and reagents to over 1,200 researchers at 750 institutions in 34 countries. BEI Resources employs a stringent authentication and characterization methodology for each strain offered through the repository. As part of its mission, BEI Resources uses the latest sequencing and bioinformatic methods to provide high-quality whole genome sequences to the research community.

The SARS-CoV-2 Wuhan-Hu-1 reference genome is 29,903 nt in length and structured similarly to closely related coronaviruses from the subgenus *Sarbecovirus*. The genomic organization includes one large gene (21,291 nt) which is cleaved into 16 nonstructural proteins (nsps) including the RNA-dependent RNA polymerase (RdRp), followed by the spike (S) protein which mediates binding to cellular receptors, an envelope (E) protein, a membrane (M) protein, a nucleocapsid (N) protein, and a variety of accessory proteins.^6^ Genomic comparison shows that the closest relative to SARS-CoV-2 is the coronavirus RaTG13, isolated from the intermediate horseshoe bat (*Rhinolophus affinis*).^7^

One of the SARS-CoV-2 accessory proteins is Protein 6 (p6), a 61 aa protein encoded by ORF6. Although its function in SARS-CoV-2 is not fully known, it bears some similarity to p6 from severe acute respiratory syndrome coronavirus (SARS-CoV), suggesting that it may play a similar role. In SARS-CoV, p6 is a 63 aa amphipathic protein that has been shown to have a variety of effects. p6 localizes to the membrane of the endoplasmic reticulum and Golgi^8^, where it serves as an antagonist of type I interferon (IFN) signaling by cytoplasmic sequestration of karyopherin α2, thus preventing the translocation of signal transducer and activator of transcription factor 1 (STAT1) into the nucleus. By preventing STAT1-activated gene expression, the cell’s ability to mount an effective antiviral response is reduced.^9^ Additionally, p6 promotes the ubiquitination and degradation of N-Myc (and STAT) interactor (Nmi), a protein that has been shown to enhance IL-2 and IFN-γ-dependent transcription.^10^ p6 also interacts directly with another IFN-associated protein, the nuclear pore complex interacting protein family, member B3 (NPIPB3).^11^ p6 has also been reported to interact with other SARS-CoV proteins, including nsp3^12^, nsp8^13^, protein 7b^12^, and protein 9b^14^.

In this work, we report the sequencing of a SARS-CoV-2 strain originally isolated from a clinical patient in Hong Kong, subsequently passaged, and then deposited with BEI Resources. Genome analysis of the deposited material shows a 27 nt in-frame deletion within ORF6, predicted to result in a 9 aa deletion from the central portion of p6. This is further predicted to result in a dramatic alteration to the predicted three-dimensional structure, possibly with significant functional implications.

## Methods and Results

### Viral Isolation and Passaging

SARS-CoV-2 Hong Kong/VM20001061/2020 was isolated from a nasopharyngeal aspirate and throat swab from 39-year-old male patient in Hong Kong on January 22, 2020.^15^ The complete genome of the SARS-CoV-2 Hong Kong/VM20001061/2020 clinical isolate (Passage 0) has been sequenced (GISAID: EPI_ISL_412028). The viral strain was isolated and passaged five times at the Hong Kong University in African green monkey (*Cercopithecus aethiops*) kidney epithelial cells (Vero E6) cells prior to its deposit with BEI Resources (Manassas, VA, USA) for production and distribution to the scientific community.

### Viral Growth and Extraction

Using the virus deposited with BEI Resources (hereinafter referred to as “HKU Passage 5”), we inoculated Vero E6 cells (ATCC^®^ CRL-1586™) using an multiplicity of infection (MOI) of 0.01, adsorbed for one hour at 37°C before adding viral growth media consisting of Eagle’s Minimum Essential Medium (ATCC^®^ 30-2003™) supplemented with 2% fetal bovine serum (ATCC^®^ 30-2020™). After 5 days of incubation at 37°C with 5% CO_2_, when roughly 75% of the cell monolayer showed cytopathic effects, the cell lysate and supernatant were harvested. This viral strain (representing Passage 6) is available from BEI Resources under the catalog number NR-52282.

Genomic RNA (gRNA) was extracted for sequencing from a preparation of cell lysate and supernatant from the Passage 6-infected cell line using QIAamp^®^ Viral RNA Mini Kit (QIAGEN). The gRNA extracted from the Passage 6 virus culture is available from BEI Resources under the catalog number NR-52388.

### Genome Sequences

The genome sequences for the viral strains analyzed are shown in **Table 1**.

**Table 1.** Viral strains and genomes used in the current work. NR-52282 (Lot# 70034432) is available from BEI Resources.

| Virus | Strain | Description | Passage # | Genome Accession |
| --- | --- | --- | --- | --- |
| SARS-CoV | Tor2 | Reference Strain | – | NC_004718.3 |
| SARS-CoV-2 | Wuhan-Hu-1 | Reference Strain | – | NC_045512.2 |
| SARS-CoV-2 | Hong Kong/VM20001061/2020 | Clinical Specimen | 0 | EPI_ISL_412028 |
| SARS-CoV-2 | Hong Kong/VM20001061/2020 | Deposited Virus* | 5 | – |
| SARS-CoV-2 | Hong Kong/VM20001061/2020 | NR-52282 | 6 | MT547814.1 |
| SARS-CoV-2 | England/CAMB-77F07/2020 | Clinical Specimen | – | EPI_ISL_441819 |
| SARS-CoV-2 | England/CAMB-722A9/2020 | Clinical Specimen | – | EPI_ISL_439593 |

\*The viral sample (*i.e.*, HKU Passage 5) deposited with BEI Resources which served as the seed for production of NR-52282.

### Viral Sequencing and Assembly

We performed sequencing library preparation using the NEBNext Ultra II RNA Library Prep Kit for Illumina (NEB Catalog #E7770). Whole-genome sequencing was performed using a v2 Micro flow cell (300-cycle) on the MiSeq^®^ system (Illumina), producing a total of 676,247 paired-end reads (1,352,494 total reads) with a length of 150 nt. Reads were then trimmed to remove low quality bases and adapter sequences, and *de novo* and reference assembly were performed. For *de novo* assembly, reads were assembled into contigs using SPAdes v3.12.0 under “RNA-seq” mode, and the resulting *de novo* contigs were filtered based on taxonomy using the One Codex Database version a76e9f6d3b23445d. Reads were then mapped to resultant *de novo* contigs using minimap2 v2.12-r836-dirty default parameters and short read preset mode `-x sr`.

Reference-based assembly was then performed by mapping the reads to the genome sequence of the clinical isolate (GISAID: EPI_ISL_412028) using minimap2 default parameters and short read preset mode `-x sr`, and mapping statistics were generated by the bamqc feature of QualiMap v2.2.1. A total of 136,746 reads mapped to the reference at a mean coverage of 640.62× (SD: 210.80×). Variants were then called, and a consensus sequence was generated using bcftools 1.10.2 with default parameters for bcftools mpileup and bcftools call, and bcftools consensus. The variants observed in NR-52282 relative to the clinical isolate genome are shown in **Table 2**. One variant, `y24034t`, was the result of an ambiguous nucleotide in the reference sequence. However, our variant calling methods report a `T` in this location with no minor variants detected (<u>**Table S1**</u>), providing strong evidence that this is the appropriate nucleotide.

**Table 2.**
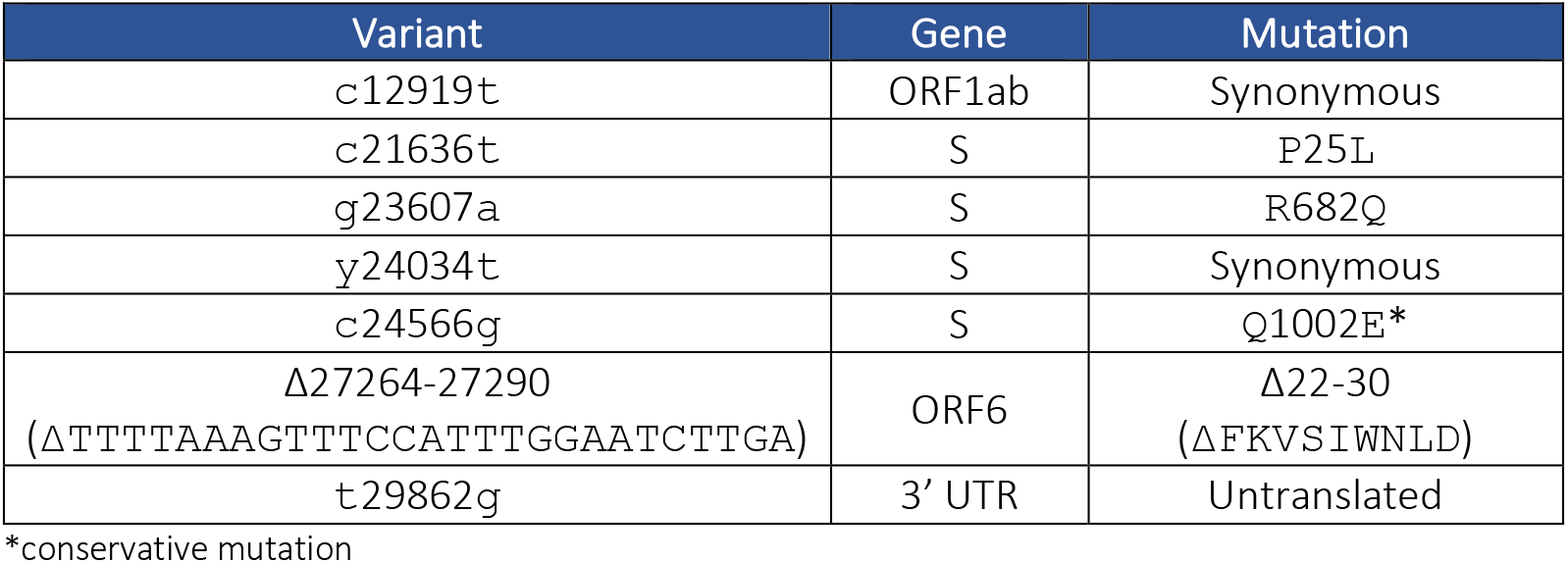
Results of variant calling between NR-52282 versus the clinical specimen genome from SARS-CoV-2 Hong Kong/VM20001061/2020 (GISAID: EPI_ISL_412028).

Of particular interest is a 27 nt deletion in ORF6, corresponding to nucleotides 27,264-27,290 in the Wuhan-Hu-1 reference genome. This is predicted to manifest as an in-frame deletion of 9 aa (`ΔFKVSIWNLD`). We refer to the predicted resultant protein as SARS-CoV-2 p6^Δ22-30^. To determine the support for this deletion, we examined the statistical support for this variant as reported by the Bcftools variant caller. Bcftools mpileup reports a quality score (PHRED-scaled) of 221, which corresponds to an error probability of 7.93 × 10^−23^, indicating that this deletion is very well supported. Furthermore, this quality is comparable to the quality scores reported for the other 6 SNPs detected (mean variant quality score = 208.67 ± 33.29), shown in **Table 2**. Furthermore, this deletion does not appear in all reads mapped to this position. Bcftools mpileup reports a maximum fraction of reads supporting this deletion as 0.706, suggesting a mixed viral population where a minority of viral genomes contain the wild-type sequence. However, an abundance of caution must be exercised in attempting to extrapolate read frequencies to the distribution of a population mix, *i.e.*, the read fraction of 0.706 does not necessarily indicate that 70.6% of the virus culture contains the deletion and the remainder does not.

To rule out possible error or bias in our chosen assembly and variant calling methods, we employed two additional orthogonal assembly methods to ascertain if these methods also recovered the deletion. We first employed the One Codex SARS-CoV-2 pipeline (https://github.com/onecodex/sars-cov-2), which also uses minimap2 to map reads to the SARS-CoV-2 reference genome (GenBank <u>MN908947.3</u>, RefSeq <u>NC_045512.2</u>), but uses iVar^16^ for viral variant calling and consensus generation. Next, we employed CDC’s Iterative Refinement Meta-Assembler (IRMA) pipeline.^17^ Briefly, IRMA uses an ambiguous consensus sequence of an RNA virus species’ genome as a seed sequence, then proceeds through multiple rounds of sample read mapping to generate a consensus sequence for that given sample. We ran a modified version of IRMA’s existing MERS module using a consensus sequence of the SARS-CoV-2 genome generated using an alignment of all existing SARS-CoV-2 genomes in GISAID as of May 13, 2020.

We aligned the One Codex consensus sequence, the IRMA consensus sequence, and our *de novo* and consensus sequences using Muscle 3.8.425. This alignment confirmed the presence of the 27 nt deletion in the One Codex and IRMA consensus sequences and the 6 SNPs detected in our *de novo* and consensus sequences. Additionally, both the One Codex pipeline and IRMA support a minority presence of the wildtype sequence in the read set. The One Codex pipeline using iVar reports the frequency of the deletion in the read set as 0.5243, while IRMA reports the frequency of the deletion in the read set as 0.0567. These results are summarized in <u>**Table S1**</u>.

In order to determine whether the deletion occurred as a result of passaging by BEI Resources, a remaining sample of the Passage 5 virus culture was also sequenced and assembled using the same methods as previous described. The alignment of the complete genomes (DNA format) of SARS-CoV-2 Wuhan-Hu-1 (wild-type), SARS-CoV-2 Hong Kong/VM20001061/2020 (GISAID: EPI_ISL_412028) from the clinical specimen, HKU Passage 5, and Passage 6 (NR-52282) is shown in <u>**Supplementary Material**</u>. We confirmed the presence of the deletion in HKU Passage 5 and noted shifts in variant frequencies between the passages (<u>**Table S2**</u>). The alignment of these genomes in the region of ORF6 is shown in **Figure 1**. The same methods were used to investigate the read-level sequencing results of the original clinical sample (Passage 0) as well as HKU Passage 2. All methods independently confirmed the absence of any Δ27,264-27,290 variants in both the original clinical specimen and HKU Passage 2 (data not shown).

**Figure 1.**
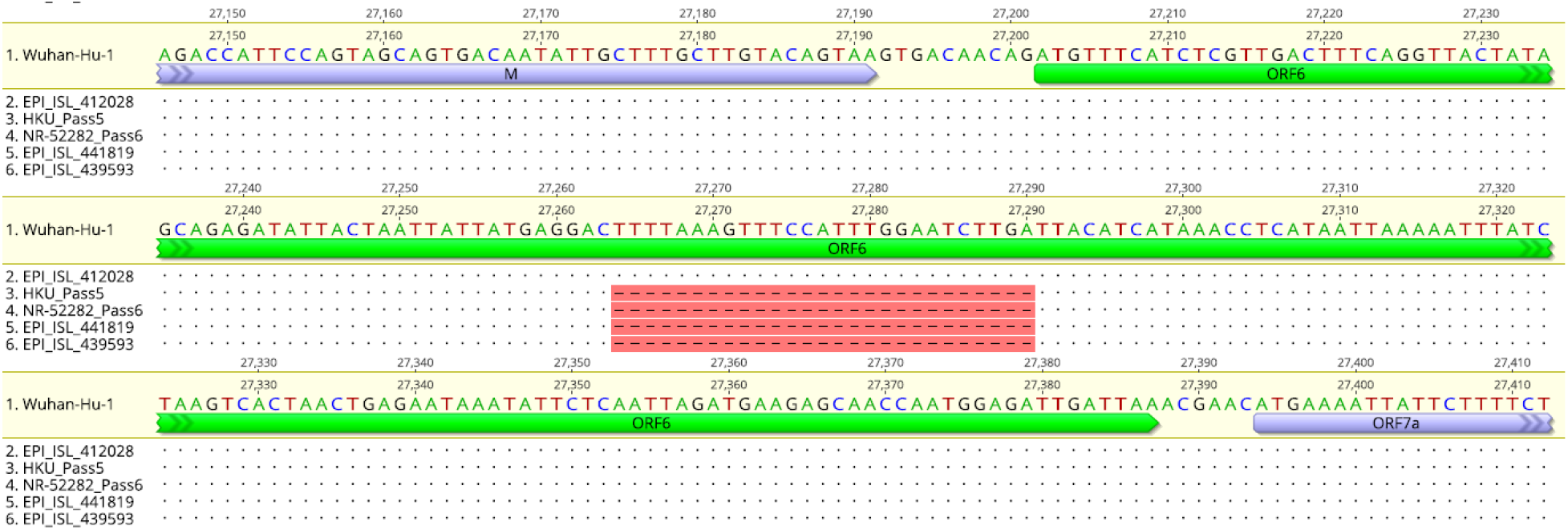
Genome alignment of SARS-CoV-2 strains in the ORF6 region. Nucleotides matching the Wuhan-Hu-1 reference strain are shown as dots. While the original clinical isolate of Hong Kong/VM20001061/2020 (EPI_ISL_412028) does not contain the 27 nt deletion, later passages of this strain (HKU Passage 5 and NR-52282 Passage 6) harbor this mutation. Also shown are two strains isolated in England (EPI_ISL_439593 and EPI_ISL_441819) which harbor the same deletion.

To determine whether similar mutations have been previously observed in SARS-CoV-2 genomes, we examined the GISAID database for mutations within ORF6. Of 24,456 strains, only two strains (hCoV-19/England/CAMB-722A9/2020: EPI_ISL_439593 and hCoV-19/England/CAMB-77F07/2020: EPI_ISL_441819) contained the same 27 nt deletion, thus encoding an identical p6^Δ22-30^. Interestingly, the GISAID metadata for these two genomes indicate that both originated from clinical specimens from the same hospital in Cambridge, United Kingdom and were collected just four days apart. These two genomes are also included in the ORF6 region alignment and the full alignment in **Figure 1** and <u>**Supplementary Material**</u>, respectively. An additional genome, hCoV-19/England/SHEF-C7788/2020 (GISAID: EPI_ISL_432847), harbors a shorter 18 nt deletion in the same region of ORF6, corresponding to nucleotides 27,273-27,290 in the Wuhan-Hu-1 reference genome (data not shown). This is predicted to manifest as an in-frame deletion of 6 aa (`ΔSIWNLD`).

### PCR Analysis and Sanger Sequencing

Primers for PCR and Sanger sequencing were designed to bind upstream and downstream of SARS-CoV-2 ORF6, thus spanning its entire sequence. We performed one-Step RT-PCR on the gRNA extracted from HKU Passage 5 and NR-52282 using the primers CoV2-ORF6 F (5’-CCATTCCAGTAGCAGTGACAATA-3’) and CoV2-ORF6 R (5’-GCTCACAAGTAGCGAGTGTTAT-3’) with Superscript III RT/Platinum Taq mix (ThermoFisher Scientific) according to the following conditions: 50°C for 15 min, 95°C for 2 min, and (95°C for 15 sec, 60°C for 30 sec) for 40 cycles. Sanger sequencing was performed on the purified PCR product using the same primers, the BigDye Terminator 3.1 system, and the ABI 3500xL Genetic Analyzer. The Sanger sequencing results clearly show high-quality basecalls throughout the entire region of the deletion, confirming that the deletion is present and not a sequencing artifact (<u>**Figure S1**</u>).

### ORF6 Protein Alignment

The p6 protein sequences from SARS-CoV Tor2, SARS-CoV-2 Wuhan-Hu-1 (wild-type), and NR-52282 were globally aligned using the Blosum62 cost matrix in Geneious Prime 2020.1.2. The alignment (**Figure 2**) clearly shows that the 27 nt deletion in the Passage 6 ORF6 manifests as a 9 aa in-frame deletion in the translated sequence.

**Figure 2.**
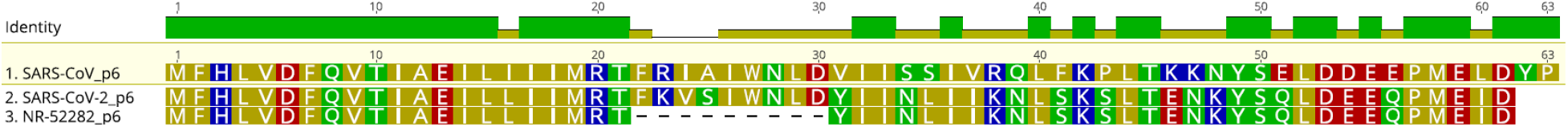
Alignment of p6 sequences from SARS-CoV Tor2, SARS-CoV-2 Wuhan-Hu-1, and SARS-CoV-2 Hong Kong/VM20001061/2020 (Passage 6; NR-52282). Residues are colored by type: nonpolar (gold), polar uncharged (green), polar acidic (red), polar basic (blue).

### Three-Dimensional Structure

In order to determine the effects that the deletion observed in ORF6 might have on its function, prediction of the folded structure was performed for p6 from SARS-CoV Tor2, SARS-CoV-2 Wuhan-Hu-1, and NR-52282 using I-TASSER.^18–20^ Three-dimensional protein structures were visualized using UCSF ChimeraX v1.0rc202005180441.^21^ Both proteins are generally structured as two α-helices separated by a flexible random coil. The predicted three-dimensional structures are shown in **Figure 3**. The SARS-CoV p6 α_1_-helix is predicted to begin at `H3` and extend to `T21`, while its α_2_-helix is predicted to begin at `W27` and extend to `Q39`; its random coil extends from `F22` to `I26` (**Figure 3A**). The SARS-CoV-2 p6 α_1_-helix is predicted to begin at `V9` and extend to `T21`, while its α_2_-helix is predicted to begin at `W27` and extend to `E46`; its random coil extends from `F22` to `I26` (**Figure 3B**). Both p6 proteins are predicted to have an extended disordered region after the second helix, from `L40` to `P63` for SARS-CoV and from `L40` to `D61` for SARS-CoV-2. In SARS-CoV, the most C-terminal portion of p6 (aa 54-63) has been found to be functionally critical in the interaction with Nmi.^10^ Because the C-terminal residues of SARS-CoV-2 p6 are highly conserved (**Figure 2**), this region may perform a similar function.

**Figure 3.**
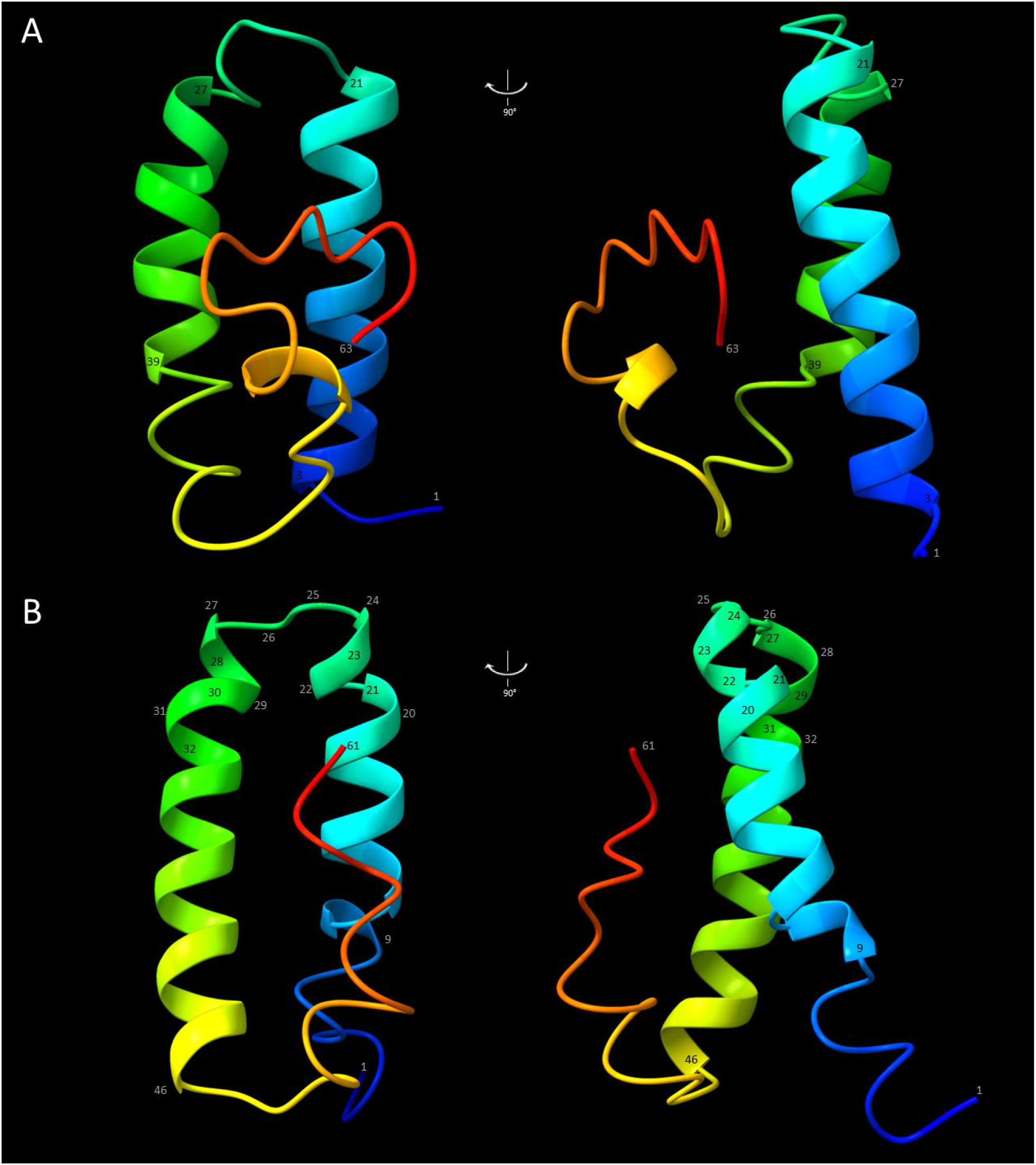
Predicted three-dimensional structures of protein 6 (p6) from A) SARS-CoV Tor2 and B) SARS-CoV-2 Wuhan-Hu-1. Structures are shown in identical perspectives, from front view (left) and side view (right). Key amino acids are labeled with their residue number.

In NR-52282, the deletion of aa 22-30 eliminates the interhelical random coil in the predicted three-dimensional structure. **Figure 4A** shows the predicted structure of SARS-CoV-2 p6 with the aa 22-30 (`FKVSIWNLD`) corresponding to the deletion from p6^Δ22-30^ highlighted in blue. In **Figure 4B**, the five amino acids before the deletion (17-21: `IIMRT`) and after (31-36: `YIINL`) are shown highlighted in red. In **Figure 4C**, the same residues (`IIMRT` and `YIINL`) are highlighted in red. These amino acids are contiguous in the predicted p6^Δ22-30^ and form the central part of a single longer α-helix.

**Figure 4.**
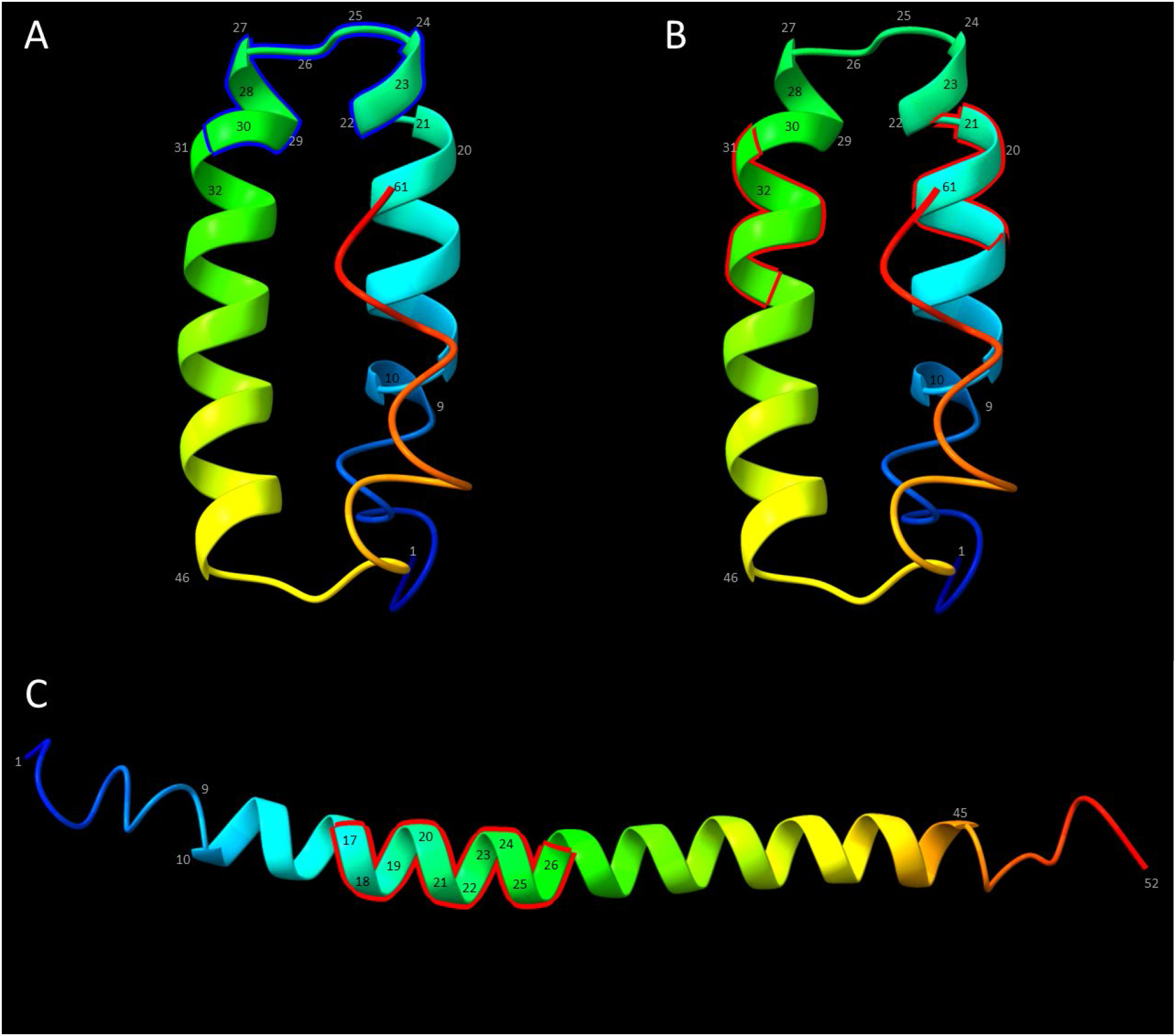
Predicted three-dimensional structure of SARS-CoV-2 p6, highlighted to illustrate A) aa 22-30 (`FKVSIWNLD`) and B) aa 17-21 (`IIMRT`) and aa 31-36 (`YIINL`). C) The predicted three-dimensional structure of SARS-CoV-2 p6^Δ22-30^. The primary consequence of the deletion is the fusion of the two α-helices into a single, longer α-helix. Highlighted are the same 10 residues highlighted in B) (`IIMRT,YIINL`), forming a contiguous sequence of residues now numbered aa 17-26.

The transmembrane potential of p6 was predicted using TMHMM2.0^22^ (<u>**Figure S2**</u>). The plots for SARS-CoV p6 and SARS-CoV-2 p6 suggest that the two α-helices and the intervening random coil may embed into the membrane with the N-terminal and C-terminal ends extending into the cytoplasm, consistent with previously hypothesized N-endo, C-endo configurations.^8,23^ However, in the NR-52282 p6, the deletion of the intervening random coil and fusion of the two α-helices into a single helix alters the predicted transmembrane orientation to N-ecto, C-endo, with aa 10-32 of the shortened 52 aa protein forming a single transmembrane helix (**Figure 5**).

**Figure 5.**
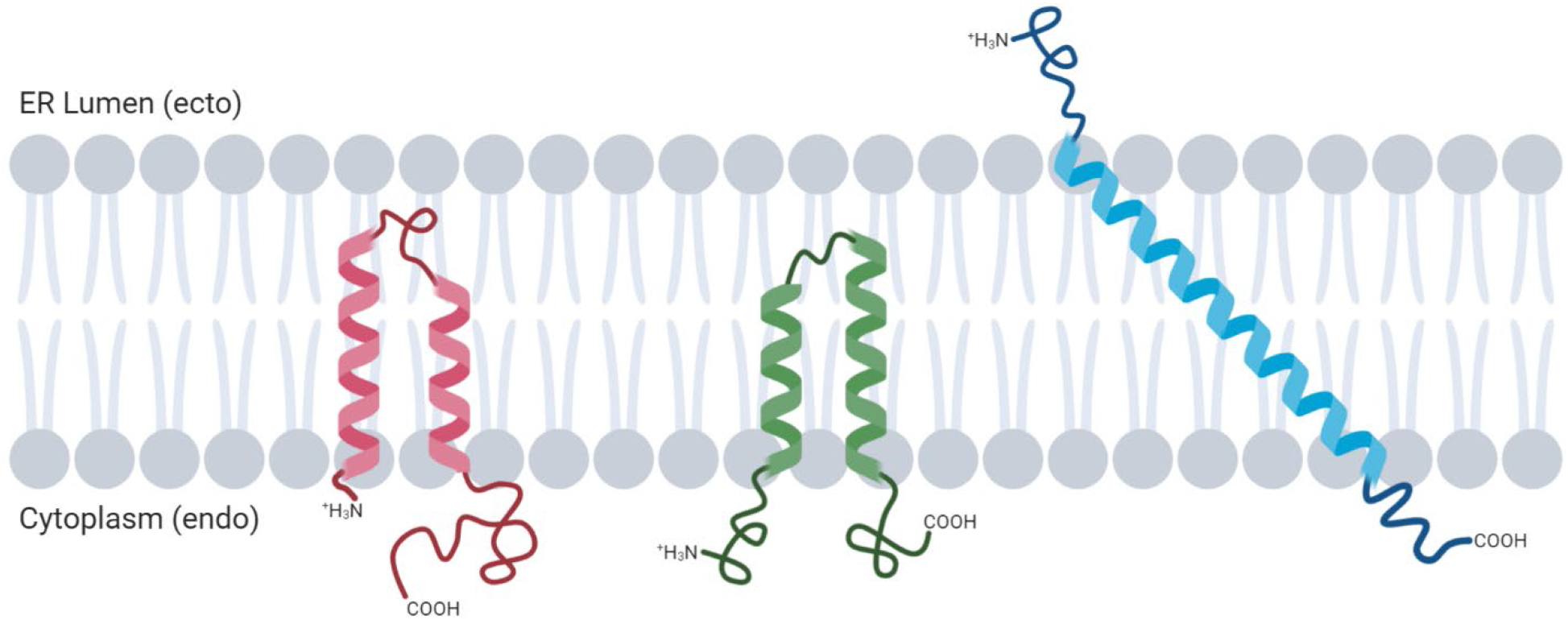
Schematic diagram of the potential transmembrane localizations of p6 from SARS-CoV Tor2 (left), SARS-CoV-2 Wuhan-Hu-1 (middle), and NR-52282 (right). Diagram created with BioRender.

The SARS-CoV p6 protein contains only two amino acids prior to the first predicted transmembrane α-helix. Because only the first two amino acids of the SARS-CoV p6 protein are predicted to extend out of the membrane, the N-terminus may not perform a function *in vivo*. However, the SARS-CoV-2 p6 contains a pre-helix sequence of eight amino acids that are predicted to extend into the cytoplasm. The specific function of the SARS-CoV-2 p6 N-terminus, if any, has not yet been determined. However, if its native function is tied to its extension into the cytoplasm, then it stands to reason that the N-endo→N-ecto shift in its localization in SARS-CoV-2 p6^Δ22-30^ (**Figure 5**) could result in the ablation of this function.

## Discussion

The Hong Kong/VM20001061/2020 strain of SARS-CoV-2 was isolated from an adult male patient in Hong Kong. The genome sequence of the clinical isolate (Passage 0) was deposited in GISAID under the accession ID EPI_ISL_412028. Prior to its acquisition by BEI Resources, the strain experienced five growth passages in Vero E6 cells. The HKU Passage 5 material acquired was used to infect Vero E6 cells; thus, the creation of the lot for distribution to the scientific community was the result of Passage 6. The 27 nt deletion (Δ27,264-27,290) seen in NR-52282 was not present in either the original clinical isolate or in the HKU Passage 2 strain^15^ but was present in the HKU Passage 5 strain received. This suggests that this deletion originated *ab initio* during Passage 3, 4, or 5. Laboratory passaging through the Vero E6 cell line may have allowed the deletion variant to become dominant. Furthermore, the p6^Δ22-30^ mutation has been reported in sequences from two UK clinical specimens, indicating that this deletion, while rare, is a naturally-occurring mutant. Further studies on the UK isolates with the p6^Δ22-30^ mutation is warranted, and data on disease severity and patient outcome could shed light on the significance of the mutant virus and the role of p6 *in vivo*.

The function of p6 in SARS-CoV-2 has not yet been fully determined. However, based on its similarity to the SARS-CoV p6, it appears likely that it plays a role as an antagonist of interferon signaling, thus assisting in the viral suppression of the innate immune system. In attempting to understand how this mutation may have propagated, it is important to consider that the Vero E6 cell line is deficient in production of interferon.^24,25^ It is possible that propagation of the virus in an IFN-deficient cell line removes the selective pressure on p6, thus allowing a mutation such as p6^Δ22-30^ to persist, whereas it may be selected against in an IFN-proficient system such as a human host.

Since SARS-CoV-2 is typically propagated using Vero E6 cells, laboratory adaptation such as that described in this work could potentially have implications if the laboratory-adapted virus produced is used for downstream studies. This also underscores the importance of limiting the number of passages in cell culture in order to maintain the wildtype properties as well the necessity of confirming by whole genome sequencing with including analysis of minor variants of each virus preparation used for research studies. Development of spontaneous mutation due to virus adaptation to both Vero and Vero E6 cells has been reported for viruses such as Ebolavirus, sometimes with altered pathogenesis.^26,27^ Similar growth adaptative mutations have also been reported with Zika virus that led to attenuation of virulence in mice.^28^ If the intent is to study the wild-type virus for basic or translational research studies, it may be useful to consider whether a more appropriate cell line could be used for SARS-CoV-2 propagation to ensure that the integrity of its native characteristics are maintained. Conversely, a laboratory-adapted strain of SARS-CoV-2 with a diminished immune evasion capacity such as the ORF6 deletion could form the basis of a vaccine candidate strain.

## Supporting information

Supplementary Alignment

## Acknowledgements

BEI Resources is funded under contract HHSN272201600013C by the National Institute of Allergy and Infectious Diseases (NIAID). The following reagents were obtained through BEI Resources, NIAID, NIH: SARS-Related Coronavirus 2, Isolate Hong Kong/VM20001061/2020, NR-52282; Genomic RNA from SARS-Related Coronavirus 2, Isolate Hong Kong/VM20001061/2020, NR-52388.

