## Supplementary Figures and Tables for "A Rare Deletion in SARS-CoV-2 ORF6 Dramatically Alters the Predicted Three-Dimensional Structure of the Resultant Protein"

### Supplementary Data

Table S1. Summary of variants called, their frequency, and reported support values for NR-52282 (Passage 6). Bcftools call variant frequency is calculated from the AD INFO field of the resultant VCF generated from bcftools mpileup using the `-a FORMAT/AD` flag where Frequency = Variant Depth / (Variant Depth + Reference Depth). Bcftools call QUAL is a Phred-scaled quality score representing probability of a correctly called variant. The iVar variant frequency is calculated from the output TSV file where Frequency = Variant Depth / (Variant Depth + Reference Depth). The iVar *p* value is the *p* value of Fisher's exact test. The IRMA variant frequency and confidence-not-machine-error is taken from the IRMA's allAlleles.txt output table, except for variant Δ27264-27290, where values are taken from IRMA's insertions.txt output table. For variant `t29862g`, iVar and IRMA values are missing; the One Codex pipeline running iVar maps against the Wuhan-Hu-1 reference genome and `G` is consistent with the reference and thus no variant is called, and IRMA's consensus sequence did not extend to that nucleotide.

| Variant | bcftools call Variant Frequency | bcftools call QUAL | iVar Variant Frequency | iVar *p* Value | IRMA Variant Frequency | IRMA Confidence Not Machine Error |
|---|---|---|---|---|---|---|
| c12919t | 0.620111732 | 222 | 0.679089027 | 8.4716E-170 | 0.689156627 | 0.999783151 |
| c21636t | 0.871657754 | 219 | 0.902155887 | 0 | 0.919921875 | 0.999791526 |
| g23607a | 0.957894737 | 228 | 0.946107784 | 0 | 0.931350114 | 0.999828475 |
| y24034t | 1 | 225 | 1 | 0 | 1 | 0.999866865 |
| c24566g | 0.725581395 | 221 | 0.736220472 | 2.1137E-196 | 0.742081448 | 0.999789404 |
| Δ27264-27290 | 0.706422 | 221 | 0.524334848 | 9.701E-212 | 0.943213296 | 0.997702313 |
| t29862g | 1 | 141 | n/a | n/a | n/a | n/a |

**Table S2.** Summary of variants called, their frequency, and reported support values for HKU Passage 5, in addition to change in variant frequency between HKU Passage 5 and NR-52282 (Passage 6). Bcftools call variant frequency is calculated from the AD INFO field of the resultant VCF generated from bcftools mpileup using the `-a FORMAT/AD` flag where Frequency = Variant Depth / (Variant Depth + Reference Depth). Bcftools call QUAL is a Phred-scaled quality score representing the probability of a correctly called variant. The iVar variant frequency is calculated from the output TSV file where Frequency = Variant Depth / (Variant Depth + Reference Depth). The iVar *p* value is the *p* value of Fisher's exact test. The IRMA variant frequency and confidence-not-machine-error is taken from the IRMA's allAlleles.txt output table, except for variant Δ27264-27290, where values are taken from IRMA's insertions.txt output table. iVar did not call `c12919t` or `c24566g` variants, while neither iVar nor IRMA called the `t29862g` variant.

| Variant | bcftools call Variant Frequency | **Bcftools call Δ Variant** Frequency | bcftools call QUAL | iVar Variant Frequency | iVar **Δ Variant** Frequency | iVar *p* Value | IRMA Variant Frequency | IRMA **Δ Variant** Frequency | IRMA Confidence Not Machine Error |
|---|---|---|---|---|---|---|---|---|---|
| c12919t | 0.266666667 | 0.353445065 | 159 | n/a | n/a | n/a | 0.3 | 0.389156627 | 0.999498055 |
| c21636t | 0.785087719 | 0.086570035 | 221 | 0.772532189 | 0.129623698 | 0 | 0.810861423 | 0.109060452 | 0.999173902 |
| g23607a | 0.814977974 | 0.142916763 | 220 | 0.795746566 | 0.150361218 | 0 | 0.79805726 | 0.133292854 | 0.998941797 |
| y24034t | 1 | 0 | 225 | 1 | 0 | 0 | 1 | 0 | 0 |
| c24566g | 0.282786885 | 0.44279451 | 180 | n/a | n/a | n/a | 0.290896646 | 0.451184802 | 0.999491672 |
| Δ27264-27290 | 0.506024096 | 0.200397904 | 222 | 0.498779343 | 0.025555505 | 0 | 0.849941838 | 0.093271458 | 0.999115048 |
| t29862g | 1 | 0 | 145 | n/a | n/a | n/a | n/a | n/a | n/a |

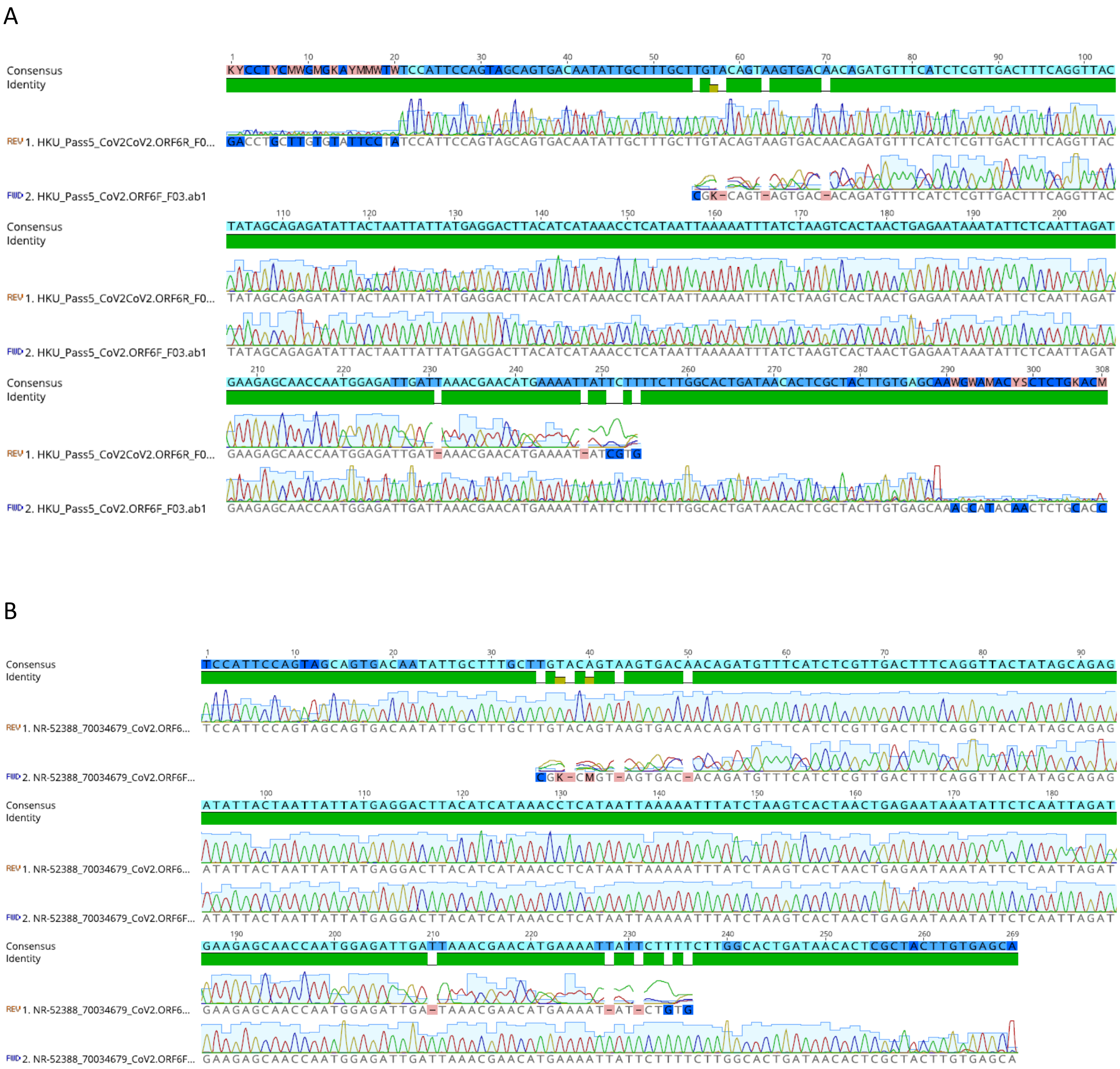


**Figure S1**. Sanger sequencing results for **A)** HKU Passage 5 and **B)** NR-52282, Passage 6 (Lot #70034679).

A

```
# SARS-CoV_p6 Length: 63
# SARS-CoV_p6 Number of predicted TMHs: 0
# SARS-CoV_p6 Exp number of AAs in TMHs: 12.55933
# SARS-CoV_p6 Exp number, first 60 AAs: 12.55933
# SARS-CoV_p6 Total prob of N-in: 0.14589
# SARS-CoV_p6 POSSIBLE N-term signal sequence
SARS-CoV_p6        TMHMM2.0     outside      1     63
```

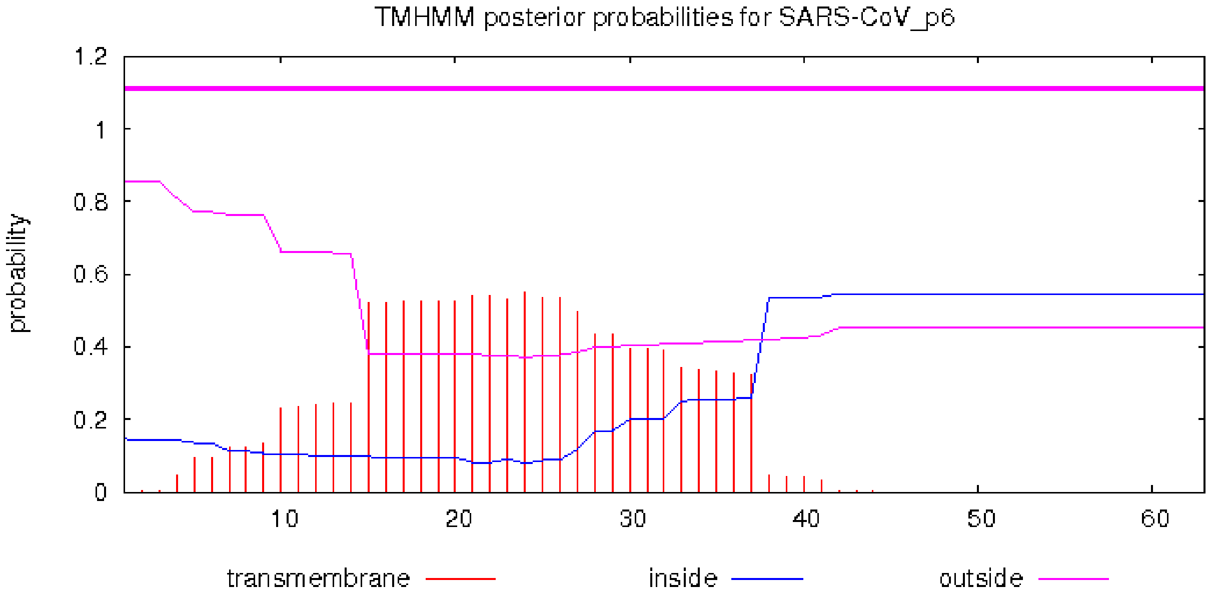


B

```
# SARS-CoV-2_p6 Length: 61
# SARS-CoV-2_p6 Number of predicted TMHs: 0
# SARS-CoV-2_p6 Exp number of AAs in TMHs: 6.09784
# SARS-CoV-2_p6 Exp number, first 60 AAs: 6.09784
# SARS-CoV-2_p6 Total prob of N-in: 0.12919
SARS-CoV-2_p6      TMHMM2.0     outside      1     61
```

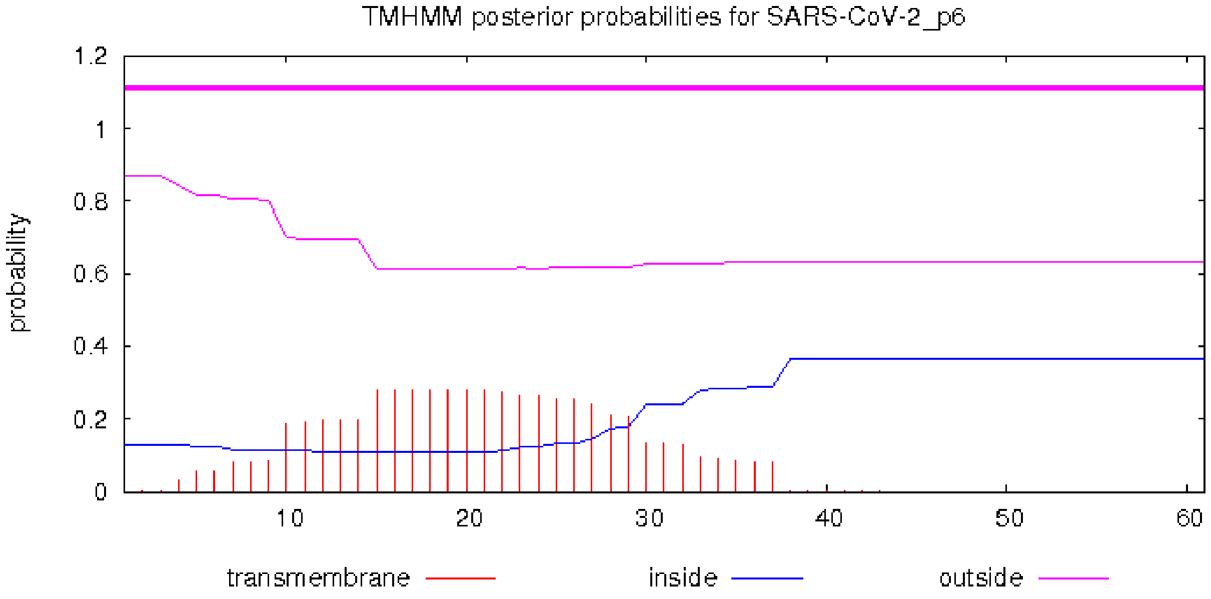


Supplementary 4

C

```
# NR-52388_p6 Length: 52
# NR-52388_p6 Number of predicted TMHs: 1
# NR-52388_p6 Exp number of AAs in TMHs: 18.68828
# NR-52388_p6 Exp number, first 60 AAs: 18.68828
# NR-52388_p6 Total prob of N-in: 0.09676
# NR-52388_p6 POSSIBLE N-term signal sequence
NR-52388_p6       TMHMM2.0    outside     1     9
NR-52388_p6       TMHMM2.0    TMhelix     10    32
NR-52388_p6       TMHMM2.0    inside      33    52
```

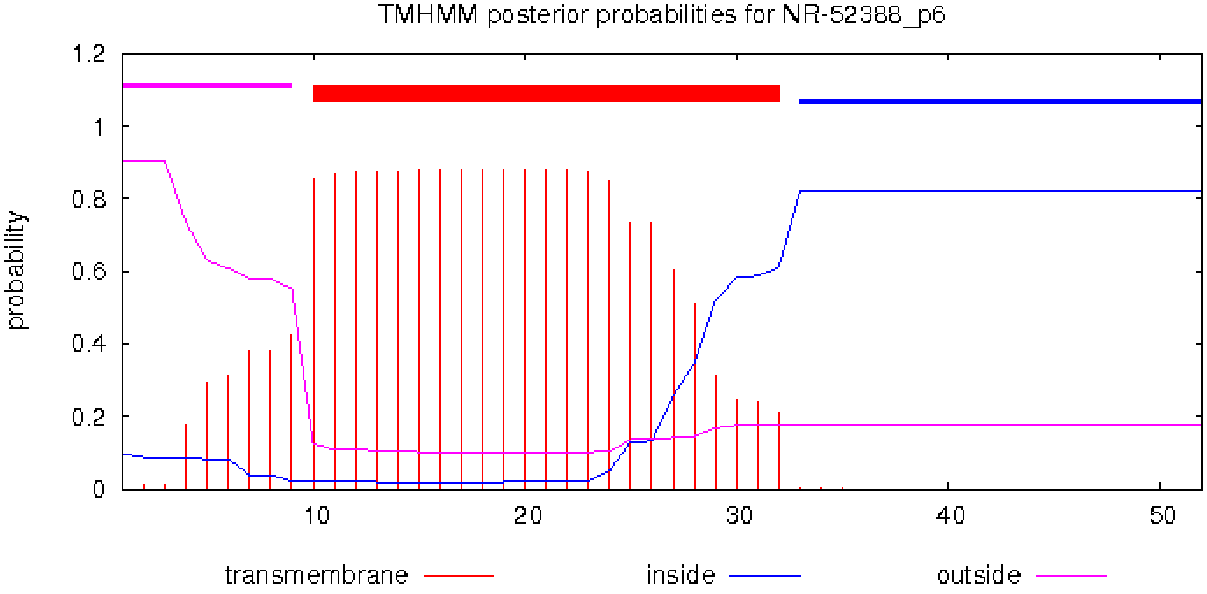


<u>Figure S2</u>. Prediction of transmembrane helices by TMHMM2.0 for protein 6 (p6) from **A)** SARS-CoV Tor2, **B)** SARS-CoV-2 Wuhan-Hu-1, and **C)** SARS-CoV-2 Hong Kong/VM20001061/2020 (Passage 6; NR-52282).
